# Spt5 phosphorylation and the Rtf1 Plus3 domain promote Rtf1 function through distinct mechanisms

**DOI:** 10.1101/2020.04.15.044131

**Authors:** Jennifer J. Chen, Jean Mbogning, Mark A. Hancock, Dorsa Majdpour, Manan Madhok, Hassan Nassour, Juliana C. Dallagnol, Viviane Pagé, David Chatenet, Jason C. Tanny

## Abstract

Rtf1 is a conserved RNA polymerase II (RNAPII) elongation factor that promotes co-transcriptional histone modification, RNAPII transcript elongation, and mRNA processing. Rtf1 function requires phosphorylation of Spt5, an essential RNAPII processivity factor. Spt5 is phosphorylated within its C-terminal domain (CTD) by cyclin-dependent kinase 9 (Cdk9), catalytic component of positive transcription elongation factor b (P-TEFb). Rtf1 recognizes phosphorylated Spt5 (pSpt5) through its Plus3 domain. Since Spt5 is a unique target of Cdk9, and Rtf1 is the only known pSpt5-binding factor, the Plus3/pSpt5 interaction is thought to be a key Cdk9-dependent event regulating RNAPII elongation. Here we dissect Rtf1 regulation by pSpt5 in the fission yeast *Schizosaccharomyces pombe*. We demonstrate that the Plus3 domain of Rtf1 (Prf1 in *S. pombe*) and pSpt5 are functionally distinct, and that they act in parallel to promote Prf1 function. This alternate Plus3 domain function involves an interface that overlaps with the pSpt5 binding site and that can interact with single-stranded nucleic acid or with the Polymerase Associated Factor (PAF) Complex *in vitro*. We further show that the C-terminal region of Prf1, which also interacts with PAF, has a similar parallel function with pSpt5. Our results elucidate unexpected complexity underlying Cdk9-dependent pathways that regulate transcription elongation.

## Introduction

Mechanisms regulating RNAPII transcription elongation are potential therapeutic targets in cancer, heart disease, and pathogenesis of HIV (1–3). Although a number of conserved positive and negative regulators of elongation have been identified, their mechanisms of action remain poorly understood (4,5). Rtf1 is a multi-functional elongation factor primarily implicated in promoting co-transcriptional histone modifications; it also has roles in RNAPII elongation and mRNA processing (4,6,7). Rtf1 is functionally linked to Cdk9, the catalytic component of P-TEFb and key driver of RNAPII elongation in all eukaryotes (4). The most extensively characterized Cdk9 targets are Rpb1, the largest subunit of RNAPII, and Spt5, an essential RNAPII processivity factor, both of which are phosphorylated on repeated amino acid motifs that comprise their C-terminal domains (CTDs)(8,9). The Spt5 CTD repeat is more variable than that of Rpb1 CTD in size and sequence, both within and between species. A related repeat motif is conserved between *S. pombe* (consensus motif: TPAWNSKS) and human [consensus motif: TP(M/L)YGS(R/Q)], in which the Thr1 residue is the Cdk9 target (10–12). The roles of the Spt5 CTD and its phosphorylation in RNAPII elongation are mostly unknown, despite the fact that it is a primary and exclusive Cdk9 target both in vitro and in vivo (8–10,13). In fact, the only established function of phosphorylated Spt5-T1 (pSpt5) is to create a binding site for Rtf1 (Prf1 in *S. pombe*)(14–16). Rtf1 recognizes pSpt5 through its conserved Plus3 domain, so named for three positively charged amino acids that are invariant among Rtf1 orthologs (17,18). The Plus3 domain is essential for the localization of Rtf1 to transcribed genes (14,17). Rtf1 also contains a highly conserved histone modification domain (HMD). The HMD directly stimulates activity of the E2 ubiquitin conjugating enzyme Rad6, leading to the mono-ubiquitylation of histone H2B (H2Bub1)(19,20). H2Bub1, in turn, directly promotes the activity of the histone H3K79 methyltransferase Dot1, as well as the Set1 histone H3K4 methyltransferase complex, and regulates chromatin dynamics during transcription (21–23). The C-terminal domain of Rtf1 interacts with the Polymerase Associated Factor (PAF) complex; this domain in human Rtf1 also stimulates RNAPII elongation *in vitro* (17,24,25). Therefore, Rtf1 Plus3 domain interaction with pSpt5 is thought to be part of a key regulatory pathway linking Cdk9 activity to co-transcriptional histone modification.

A crystal structure of the Plus3 domain in complex with phosphorylated Spt5 CTD has provided a high-resolution view of this interaction, and mutations that eliminate or decrease the interaction between the Plus3 domain and pSpt5 abrogate Rtf1 association with transcribed genes *in vivo* (15). Similarly, Spt5 CTD mutations that eliminate the Cdk9-dependent phosphorylation site also prevent the association of Rtf1 with chromatin and diminish H2Bub1 levels (14,16,26,27), consistent with pSpt5 recognition by Plus3 domain playing a central role in Rtf1 function. However, the Plus3 domain has also been shown to have other functions. For example, Plus3 contains a subdomain with structural similarity to the nucleic acid-binding PAZ domains found in Argonaute family proteins (18,28). The Plus3 domain has been shown to interact with single-stranded DNA (ssDNA) *in vitro*. The physiological significance of the nucleic acid interaction is not understood, nor is the relationship between pSpt5 binding and nucleic acid binding. Previous biochemical studies argue that these two functions are likely to be separable, although this has not been formally tested (18,28). In the fission yeast *S. pombe*, genetic ablation of H2Bub1 or of the Rtf1 ortholog Prf1 cause cell division and morphology phenotypes that are not caused by Spt5-T1 mutations (13,16,29,30). These data call into question the idea of a simple, linear pathway connecting Cdk9 activity to H2Bub1 through Plus3 domain binding to pSpt5.

We have used the model eukaryote *S. pombe* to evaluate the physiological significance of the putative Cdk9-Spt5-Prf1 pathway. Surprisingly, our data suggests that both pSpt5 and the Prf1 Plus3 domain act independently to mediate Prf1 function in elongation. The additional Plus3 domain interaction involves an interface that overlaps the pSpt5 binding site, is necessary for Prf1 chromatin association, and shares function with a C-terminal region of Prf1 that interacts with the PAF complex. Our results suggest that recruitment of Prf1/Rtf1 to sites of transcription involves multiple interactions that are modulated both directly and indirectly by Cdk9-dependent Spt5 phosphorylation.

## Materials and Methods

### Yeast strains

*S. pombe* strains used in this study are listed in Table S1. All genetic manipulations were conducted using standard techniques as previously described (31). Standard YES media (5g/L yeast extract, 30g/L D-glucose, 250mg/L of each histidine, leucine, adenine, and uracil) and 30◦C was used for the growth of all liquid cultures.

To integrate C-terminal truncation mutations into the chromosomal *prf1+* locus, primers were designed to amplify the C-terminal TAP tag from pJT9 (pFA6a-kanMX6-CTAP2) as described (32). PCR products were transformed into competent JT204 *S. pombe* cells as described (33). To integrate Plus3 domain point mutations into the chromosomal *prf1+* locus, EagI-XhoI digests of plasmids pJT161, pJT162, or pJT163 (described below) were transformed into competent JT204 cells as described above. Positive transformants were verified by sequencing and western blotting.

### Plasmids

Plasmids used in this study are listed in Table S2. Full-length Prf1, the N-terminal region (amino acids 1-213), the Plus3 domain (amino acids 214-345), and the C-terminal region (amino acids 346-563) were PCR amplified from *S. pombe* cDNA and cloned into pGEX-6P-1. For the full-length protein, a C-terminal 6xhistidine tag was introduced by PCR. Plus3 domain point mutations were introduced using the Phusion Site-Directed Mutagenesis kit (ThermoFisher Scientific) and verified by sequencing.

To integrate mutations into the chromosomal *prf1*^*+*^ locus, a ~4.5 kilobase region spanning the locus and including ~250 base pairs of 5’ and 3’ homology was PCR amplified from strain JT202 (*prf1-TAP::kanMX6*) and cloned into pGEX-6P-1 to create pJT150. Plus3 domain mutations were introduced by site-directed mutagenesis as described above. Wild-type and mutant *prf1-TAP::kanMX6* constructs were verified by sequencing.

### Expression and purification of recombinant Prf1

GST-fusion proteins were expressed in *E. coli* BL21. Log-phase cultures (500 mL) were induced with 1mM isopropyl-ß-D-1-thiogalactopyranoside (IPTG) and grown at 16°C for 12-16 hours. The cells were then harvested by centrifugation and resuspended in 25 mL of lysis buffer (20mM Tris [pH 7.5], 200mM NaCl, 20% glycerol, 1mM ethylenediamine tetra acetic acid (EDTA), 1mM dithiothreitol (DTT), 1mM phenylmethylsulfonyl fluoride (PMSF), 1 mM benzamidine, protease inhibitor cocktail (Roche Applied Sciences)) with 2.5 mg of lysozyme. After 30 mins on ice, the cell extract concentration was adjusted to 350mM of NaCl and 0.5% Triton X-100 and then sonicated with the Misonix Sonicator 3000 (30 s ON/OFF for 14 rounds, output 5.0). All subsequent steps were conducted at 4◦C. The suspension was centrifuged for 10 minutes at 25,000g and the lysate supernatant was incubated for 3 hours with 1 mL of glutathione sepharose beads (GE Healthcare) pre-washed in lysis buffer. The beads were collected, transferred to a small column (Bio-Rad), and washed with 20 mL of wash buffer (20mM Tris [pH 7.5], 350mM NaCl, 20% glycerol, 1mM EDTA, 0.1% Triton X-100, 1mM DTT, 1mM PMSF, 1mM benzamidine). Beads were eluted stepwise with 10×0.5 mL elution buffer (20mM Tris [pH 7.5], 350mM NaCl, 20% glycerol, 1mM EDTA, 100mM reduced glutathione, 1mM DTT, 1mM PMSF, 1mM benzamidine). Peak fractions were pooled and dialyzed overnight against 2L of dialysis buffer (20mM HEPES [pH 7.6], 20% glycerol, 0.15M KOAc, 10mM Mg(OAc)_2_, 1mM EDTA, 1mM DTT). For full-length Prf1, a second purification step was performed before dialysis. Briefly, eluates were supplemented with imidazole at a final concentration of 10mM and incubated for 2 hours with 200 µL of Ni-NTA agarose beads (Qiagen) that were prewashed in buffer C (20mM HEPES [pH 7.6], 150mM KCl, 5% glycerol, 10mM imidazole, 0.1% NP-40, 1mM PMSF, 1mM β-mercaptoethanol). The beads were collected, washed four times with 1 mL of buffer C, and eluted into 1 mL elution buffer 2 (20mM Tris [pH 7.5], 150mM KCl, 5% glycerol, 200mM Imidazole, 0.1% NP-40, 1mM PMSF, 1mM β-Mercaptoethanol). The eluate was then dialyzed overnight as described above.

### Immobilized Peptide Binding Assays

Spt5-CTD peptides (16) were synthesized as described (34). 10 µg of either phosphorylated or unphosphorylated peptide were immobilized on 15 µL of pre-washed streptavidin Dynabeads (Invitrogen) in 200 µL 1X PBS. After a 3-hour incubation at room temperature on a rocking platform, beads were collected on a magnet and washed twice with wash buffer (20mM HEPES [pH 7.6], 5% glycerol, 0.1% Triton X-100, 1mM EDTA, 350mM KOAc, 10mM β-glycerophosphate, 1mM PMSF). 50 ng of purified protein was added to beads and the volume made to 200 µL with binding buffer (20mM HEPES [pH 7.6], 0.1% Triton X-100, 50mM KOAc, 10mM β-glycerophosphate, 1mM PMSF, 1mg/mL BSA). The reaction was incubated at 4◦C for 1 hour with rocking. The beads were collected, washed four times with 1mL of wash buffer, and resuspended in 20 µL of 1X SDS sample buffer. All samples (5% input and 50% beads) were boiled at 95◦C for 2 minutes, centrifuged, and analyzed by SDS-PAGE and immunoblotting.

### GST Pulldowns

GST-fusion proteins and purified factors were added in equimolar amounts (approximately 20nM) in a 200µL binding reaction containing 20mM HEPES [pH 7.6], 0.1% Triton X-100, 50mM KOAc, 1mM PMSF, 10mM β-glycero-3-phosphate, 0.1 mg/mL BSA. Binding reactions were incubated for 2 hours at 4◦C on a rocking platform. GST-fusions were recovered by addition of 25 µL of glutathione sepharose beads (prewashed twice with 400 µL of binding buffer) and incubation for a further 1 hour at 4◦C with rocking. The beads were collected and then washed four times with wash buffer (20mM HEPES [pH 7.6], 5% glycerol, 0.1% Triton X-100, 1mM EDTA, 350mM KOAc, 10mM β-glycerophosphate, 1mM PMSF). The beads were then resuspended with 25µL of 1X SDS sample buffer. All samples (5% input and 50% beads) were boiled at 95◦C for 2 minutes, centrifuged, and analyzed by SDS-PAGE and immunoblotting.

### Immunoblotting

Whole-cell extracts prepared in trichloroacetic acid or purified proteins were analyzed by SDS-PAGE and immunoblotting as described previously (13). The following commercial antibodies were used: TAP tag (ThermoFisher Scientific #PICAB1001), Rpb1 (8WG16; Covance #MMS-126R-200), histone H3 (Abcam #ab1791), ubiquityl-histone H2B (Millipore #05-1312-I), GST-tag (ThermoFisher Scientific #8-326), HIS-tag (Sigma-Aldrich H1029), and Streptactin-HRP (ThermoFisher Scientific #21130). Alpha-tubulin antibody (TAT-1) was provided by Dr. Keith Gull (35). Images were acquired on Amersham Imager 600 (GE Healthcare) or on film. Images were processed using ImageJ software for quantification.

### Electrophoretic Mobility Shift Assays

Reactions contained 0.1µM of FITC-labeled deoxynucleotide probe [5’-CCGCCCCGCC-(T)_10_-CCCGCCGCCC-FITC], 10mM Tris [pH 7.5], 10% glycerol, 100mM NaCl, 0.1 mg/mL BSA, and 0.1-0.5µM recombinant GST-Plus3 protein in a final volume of 20 µL. For reactions containing the “bubble” probe the probe above was first hybridized to 5’-GGGCGGCGGG-(T)_10_-GGCGGGGCGG. After a 20-minute incubation on ice, reactions were briefly centrifuged and loaded on native 5% polyacrylamide gels. Gels were prepared and run in 0.5X Tris-borate-EDTA and run at 100 V for 1 hour at 4°C. Images were acquired on the Typhoon Trio Variable Mode Imager system (GE Healthcare). Rpb1 peptides used in competition experiments were synthesized as described (33) with sequence biotin-PSYSPTSPSYSPTSPS (unphos) or biotin-PSYSPTS*PSYS*PTSPS (phos; asterisks follow phosphoserines).

### TAP-tagged Protein Purification

Tpr1-TAP was purified from whole cell extracts as described previously (16,36). 10 µL of the purified material was analyzed by SDS-PAGE alongside BSA standards followed by Coomassie or silver staining.

### Fluorescence Microscopy

Diamino-phenylindole (DAPI) and calcofluor staining was conducted as previously described with minor changes (13). Cells were viewed using a Leica DM 5000b microscope with Lumenera’s Infinity 3-1UR camera at 40X objective. Images were processed and cells were counted using ImageJ software. Phenotypes scored were unseparated chains of cells (with septa in between each nuclei) and “twinned” septa (multiple septa separating two nucleic) (13,16). Each strain was scored three times based on images of >100 cells.

### Chromatin Immunoprecipitation (ChIP)

TAP-ChIP was conducted as previously described (37) with some modifications. ChIP experiments were normalized to spiked-in *S. cerevisiae* chromatin (prepared from strain JTY41 (genotype *MAT*a *his3Δ1 leu2Δ0 LYS2 ura3Δ0 RPB1-TAP::kanMX6*)) unless otherwise indicated. *S. pombe* chromatin was prepared as described using 1.5×10^7^ cells per crosslinked sample in a final volume of 1mL. *S. cerevisiae* chromatin was prepared using the same protocol but with 3.0×10^7^ cells per crosslinked sample. 50µL of *S. cerevisiae* spike-in chromatin was added to each *S. pombe* chromatin sample prior to conducting the immunoprecipitation (IP) step. A 100 µL sample of input was then taken from each sample. The IP was conducted by adding 20µL of IgG sepharose beads (GE Healthcare) to each IP sample for a 4-hour incubation at 4◦C. Following the IP, the beads were eluted and then washed with 150µL of TE. For the DNA purification, all samples were incubated with 0.5 µL of RNase A (10 mg/mL) and 1 µL of glycogen (20 mg/mL) for an hour at 37◦C, followed by 1.25 µL Proteinase K (20 mg/mL) for 3 hours at 37◦C. Samples were then extracted with 250 µL of phenol:chloroform:isoamyl alcohol and 250 µL chloroform as described. Primers for qPCR analysis are listed in Table S3. A primer pair in the coding region of the *S. cerevisiae PMA1* gene was used for normalization.

## Results

### Functional divergence of Prf1 Plus3 domain and phosphorylated Spt5

To examine the physiological significance of pSpt5 binding by the Plus3 domain in *S. pombe*, we introduced a point mutation at Arg227 of *prf1*^+^ that is predicted to disrupt the pSpt5 binding pocket (R227A; Figure 1A) (15,18). We verified the effect on pSpt5 interaction using immobilized peptide pulldowns. Biotinylated peptides corresponding to either unmodified or phosphorylated Spt5 CTD repeats were immobilized on streptavidin beads and incubated with purified recombinant Plus3 domain, and bound proteins were analyzed by immunoblot. Quantification of the immunoblot signals showed that wild-type Plus3 preferentially bound to the pSpt5 peptide compared to the unmodified peptide. This preference was completely abrogated by the R227A mutation, as expected (Figures 1B and S1A). We also verified that the HMD and C-terminal regions of Prf1 did not contribute to pSpt5 binding (Figure S1A and S1B).

**Figure 1.**
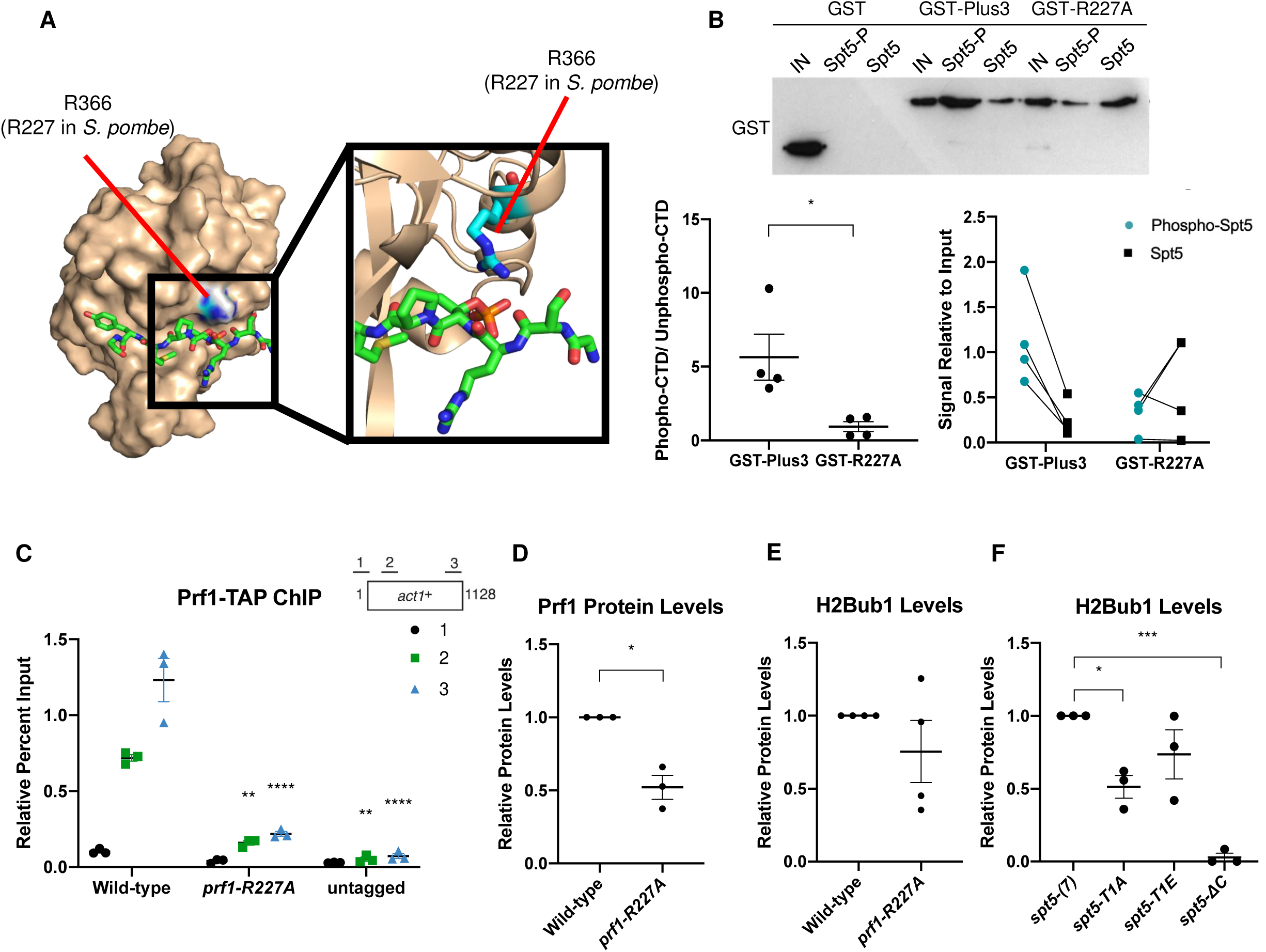
The *prf1-R227A* mutation abolishes pSpt5-binding and chromatin association, but preserves Prf1 function. **(A)** Pymol illustration mapping the location of Prf1 R227 on the crystal structure of the human Plus3 domain in complex with a pSpt5 peptide (PDB 4L1U). R366 is the equivalent position in the human protein (15). **(B)** Immobilized peptide pulldowns with the indicated Spt5 CTD peptides and the indicated recombinant GST fusion proteins. Binding reactions were analyzed by SDS-PAGE and immunoblotting with GST antibody. Top: Representative GST immunoblot. “IN” denotes a 10% input. Bottom left: Quantification of ratio between bound signal of phosphorylated Spt5 CTD and unphosphorylated Spt5 CTD peptides. Error bars denote standard error of the mean from 4 independent experiments. * p≤ 0.05; two-sided t-test. Bottom right: Quantification of bound signal relative to input for each of the 4 independent experiments. Lines between phosphorylated Spt5 CTD and unphosphorylated Spt5 CTD indicate corresponding signals within each experiment. **(C)** TAP-tag ChIP was performed on the indicated strains and quantified with qPCR using the indicated primers in *act1*^*+*^; % IP values were normalized using a primer pair in the *S. cerevisiae PMA1* gene. Length of gene (in base pairs) and position of PCR amplicons shown in diagram at the top. Error bars denote standard error of the mean from 3 independent experiments. A two-way ANOVA was conducted followed by two-sided t-tests with Bonferroni correction between each strain and wild-type within a specific primer pair. ** p≤ 0.01, **** p≤ 0.0001. **(D)** Quantification of immunoblots analyzing Prf1-TAP protein levels normalized to tubulin and then wild-type for the *prf1-R227A* strain. **(E)** Quantification of H2Bub1 levels normalized to total H3 levels and then wild-type for the *prf1-R227A* strain. **(F)** Quantifications of H2Bub1 levels normalized to total H3 levels in *spt5* mutant strains. *spt5-(7)* levels were set to 1. For (D)-(F), error bars denote standard error of the mean from 3 independent experiments. A one-sample two-sided t-test was conducted between each strain and its relative normalized wild-type. * p≤ 0.05, *** p≤ 0.001.

To examine the impact of pSpt5 binding on Prf1 function *in vivo*, we introduced the R227A mutation into the endogenous *prf1*^*+*^ locus and compared it to the effect of mutations in the Spt5 CTD that abolish all of the Cdk9-dependent phosphorylation sites. Spt5 CTD mutations (T1A or T1E) were engineered in the context of a truncated, 7-repeat *spt5*^*+*^ CTD domain whose function is comparable to wild-type [*spt5(7)*] (16,29,38). We also analyzed the *spt5-ΔC* mutant in which the entire CTD is deleted. In chromatin immunoprecipitation (ChIP)-qPCR assays, Prf1-R227A recruitment to transcribed genes was significantly decreased (up to 5-fold) throughout gene bodies compared to wild-type to levels close to those obtained in the untagged control (Figures 1C and S2). A comparable effect on Prf1 chromatin association was elicited by *spt5-T1A* and *spt5-T1E* mutants (16). The *prf1-R227A* mutation caused a more modest, two-fold reduction in Prf1 protein levels, which argues that the reduced chromatin occupancy reflects an impaired interaction (Figures 1D and S3). Despite the strong effects of Plus3 or Spt5 phosphorylation mutations on Prf1 chromatin occupancy, we observed relatively modest effects on H2Bub1 levels. Immunoblotting of whole-cell extracts indicated that H2Bub1 levels were not significantly affected by *prf1-R227A* and were reduced two-fold by *spt5-T1A;* a complete loss was observed in the *spt5-ΔC* mutant (Figures 1E, 1F, and S3). These results suggest that the Plus3 domain/pSpt5 interaction is necessary for Prf1 chromatin binding but is only partially required for mediating Prf1 function and the effect of Cdk9 activity on H2Bub1.

Genetic ablation of H2Bub1 (*htb-K119R*) or of Prf1 (*prf1Δ*) lead to cell division defects (13,16,30). These include unseparated chains of cells with division septa in between each nuclei, and “twinned” septa (multiple septa separating two nuclei) (13,16). We examined *prf1-R227A* cells stained with DAPI and calcofluor by fluorescence microscopy and saw no differences from wild-type (Figure 2A; quantified in Figure 2B). The *spt5-T1A* mutant was also similar to wild-type in these assays, consistent with the partial requirement of the Plus3 domain/pSpt5 interaction for H2Bub1 formation (Figure 2B) (29).

**Figure 2.**
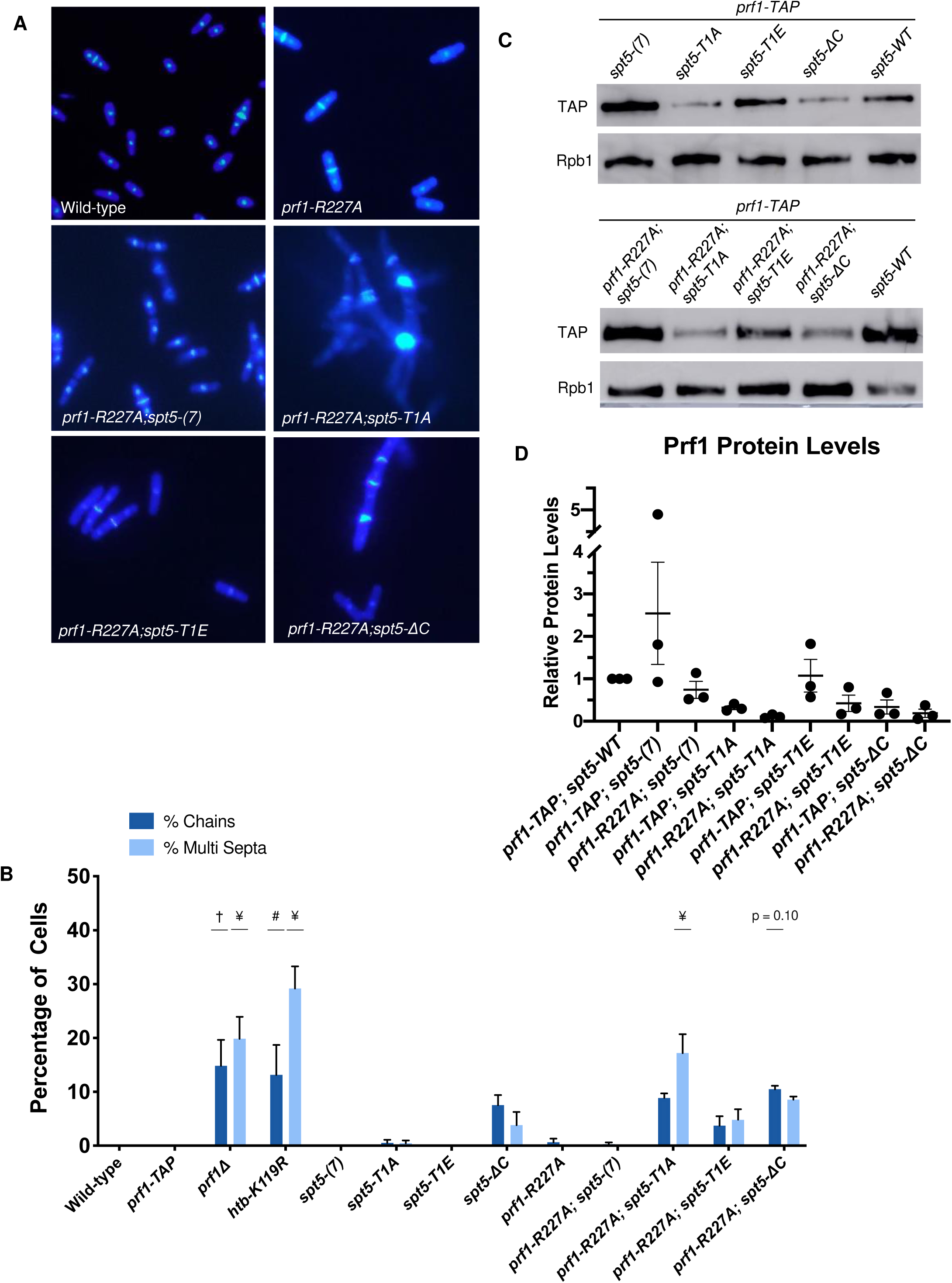
The Plus3 domain and pSpt5 function in parallel pathways. **(A)** The indicated strains were stained with DAPI and calcofluor and visualized by fluorescence microscopy. **(B)** Quantification of septation defects in indicated strains normalized to the number of septated cells counted in each indicated strain. Error bars represent standard error of the mean from 3 independent experiments; at least 100 cells were counted for each strain per experiment. A one-way ANOVA was conducted across all strains followed by two-sided t-tests with Bonferroni correction between each strain and the wild-type *prf1-TAP* strain, for each specific morphology defect. # p≤ 0.01, † p≤ 0.001, ¥ p≤ 0.0001. **(C)** Immunoblots of whole cell extracts from the indicated strains. Antibodies are indicated on the left. **(D)** Quantification of Prf1-TAP protein levels in *prf1-R227A spt5* double mutant strains and *spt5* single mutant strains. Ratios of TAP/Rpb1 signals for each sample were normalized to that in *prf1-TAP spt5*^*+*^. Error bars denote standard error of the mean from 3 independent experiments. A one-way ANOVA was conducted across all *prf1* strains within a *spt5* background followed by two-sided t-tests with Bonferroni correction between each *prf1* mutant strain and the wild-type *prf1-TAP* strain in the same *spt5* background.

We also created double mutants harboring *prf1-R227A* in combination with each of the *spt5* mutations. Given that the R227A mutation lies within a well-characterized binding site for pSpt5, we anticipated that any phenotypic effects would be due solely to loss of pSpt5 binding, and thus we predicted an epistatic relationship between *prf1-R227A* and *spt5-T1A*. Surprisingly, when the *prf1-R227A* mutation was combined with the *spt5-T1A*, the resulting double mutants displayed a significant increase in the cell division phenotypes characteristic of *prf1Δ* and *htb-K119R* cells (Figures 2A and 2B; the overall septation indices for these strains are shown in Figure S4). This indicated a synthetic, rather than epistatic relationship between the mutations, which argues that they affect different pathways. No phenotype was observed in strains with *prf1-R227A* in combination with the *spt5(7)* allele. The modest synthetic effects observed in combination with the *spt5-T1E* or *spt5-ΔC* alleles were not statistically significant, indicating that the synthetic effects were specifically related to loss of pSpt5.

We performed immunoblots to monitor Prf1 protein levels in single and double mutant strains. Interestingly, the *spt5-T1A* and *spt5-ΔC* mutations, but not *spt5-T1E*, resulted in decreased Prf1 protein levels in the wild-type *prf1-TAP* strain, supporting a direct or indirect functional link between pSpt5 and Prf1. However, introduction of *prf1-R227A* into these strains did not significantly reduce Prf1 levels relative to those in wild-type *prf1-TAP* (Figures 2C and 2D). We conclude that the synthetic phenotypes observed in *prf1-R227A spt5-T1A* double mutants are due to an impaired Prf1 interaction distinct from that with pSpt5.

The *prf1Δ* and *htb-K119R* mutants are sensitive to thiabendazole (TBZ), a microtubule-destabilizing agent that perturbs mitotic chromosome segregation, and methyl methanesulfonate (MMS), a DNA-damaging agent (39–41)(Figure S5). To determine whether these phenotypes were subject to similar synthetic effects, we assessed growth of *prf1-R227A, spt5* CTD single mutants, and double mutants in the presence of TBZ or MMS. The double mutant strain with *spt5-T1A*, but not with *spt5(7)* or *spt5-T1E*, showed a marked decrease in growth on the control media compared to either single mutant, consistent with the observed cell division phenotypes (Figure S5). In the presence of either TBZ or MMS, growth of this double mutant was specifically suppressed, whereas no effect on growth was observed for either single mutant or the other double mutant combinations. The *spt5-ΔC* mutant was sensitive to TBZ and MMS on its own, and this sensitivity was enhanced in combination with *prf1-R227A*. Together, these synthetic phenotypes establish an additional function for the pSpt5-binding surface of the Plus3 domain, as well as a Plus3-independent function for pSpt5.

### Nucleic acid binding activity of the Plus3 domain is required for Prf1 recruitment to chromatin and is competitive with pSpt5 binding

We investigated whether the additional function of the Plus3 domain could be attributed to its ability to bind nucleic acids. We first used electrophoretic mobility shift assays (EMSA) to assess binding of recombinant Prf1 Plus3 domain to a fluorescently labeled ssDNA probe (30 deoxynucleotides in length). In the presence of increasing amounts of Plus3 domain, the intensity of the band corresponding to the free probe diminished and intensity of a diffuse band close to the well increased; we also noted a general increase in signal intensity throughout the lane (Figure 3A; see arrow). This pattern likely reflects formation of heterogeneous protein-nucleic acid complexes, similar to what was previously observed for the Plus3 domain from human Rtf1 (18). Prf1 Plus3 domain also bound a “bubble” DNA substrate (a double-stranded probe with a central region of non-complementarity designed to mimic the transcription bubble) (Figure 3A). Competition experiments showed no apparent binding to dsDNA, but indicated that RNA competes for binding to labeled ssDNA probe just as or more effectively than ssDNA (Figure S6A-C). Thus, nucleic acid binding is a conserved property of the Plus3 domain.

**Figure 3.**
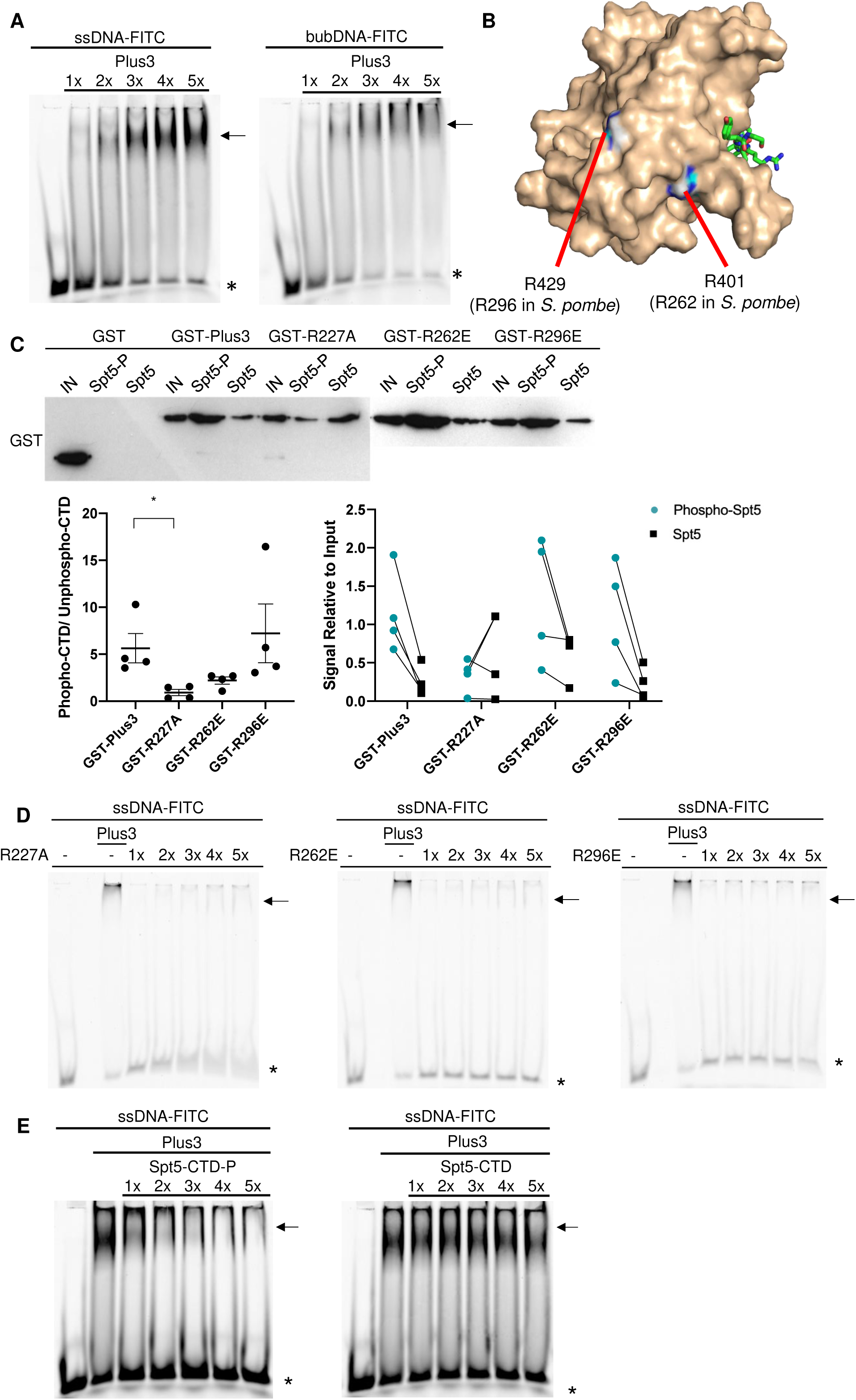
Nucleic acid binding activity of the Plus3 domain is mutually exclusive with pSpt5 binding. **(A)** EMSAs containing a FITC-labelled ssDNA (left) or “bubble” (bubDNA; right) DNA probe and a 1x to 5x molar equivalent of the Prf1 Plus3 domain. The predominant shifted band is denoted with an arrow. * indicates the free probe. All experiments were repeated at least 3 times and representative images are shown. **(B)** Pymol illustration mapping the location of Prf1 R262 and R296 on the human Plus3 domain/pSpt5 crystal structure (PDB 4L1U). Conservation of the equivalent positions in the human protein is indicated (15). **(C)** Immobilized peptide pulldowns with the indicated Spt5 CTD peptides and the indicated recombinant GST fusion proteins. Binding reactions were analyzed by SDS-PAGE and immunoblotting with GST antibody. Left: Representative GST immunoblot (blot for GST, GST-Plus3, GST-R227A is reproduced from Figure 1B). “IN” denotes a 10% input. The left half of this blot is identical to Figure 1B. Middle: Quantification of ratio between bound signal of phosphorylated Spt5 CTD and unphosphorylated Spt5 CTD peptides. A two-sided t-test was conducted between the each Plus3 mutant and the Plus3 wild-type signal ratios. Error bars denote standard error of the mean from 4 independent experiments. * p≤ 0.05. Right: Quantification of bound signal relative to input for each of the 4 independent experiments. Lines between phosphorylated Spt5 CTD and unphosphorylated Spt5 CTD indicate corresponding signals within each experiment. **(D)** EMSAs containing a FITC-labelled ssDNA probe and a 1x to 5x molar equivalent of the indicated Prf1 Plus3 domain. Lane marked “Plus3” contains wild-type Plus3 domain at 1x concentration. **(E)** Competition experiments containing ssDNA probe, a 1x molar equivalent of Prf1 Plus3 domain, and Spt5-CTD peptide (either phosphorylated or unphosphorylated) added at 1x to 5x molar ratio to probe. For (D) and (E) experiments were repeated at least 3 independent times and representative images are shown.

Previous studies showed that residues in the predicted PAZ subdomain, distant from the pSpt5 binding pocket, were critical for ssDNA binding (15,18). We substituted two equivalent positions in the Prf1 Plus3 domain-Arg262 and Arg296-with glutamates (Figure 3B). These substitutions had minimal effect on interaction with pSpt5 in immobilized peptide pulldowns, although the R262E mutation decreased (but did not eliminate) phospho-binding preference (Figure 3C). As was found for human Plus3, both mutations abolished the nucleic acid-binding of Prf1 Plus3 (Figure 3D). Interestingly, the R227A mutation also dramatically decreased the Plus3 domain’s nucleic acid binding activity, although some mobility shift could be discerned at higher protein concentrations (Figure 3D). This suggests that pSpt5-binding and nucleic acid-binding functions may reside in overlapping regions of the Plus3 domain. To confirm this, we performed EMSA assays with wild-type Plus3 domain in the presence of either phosphorylated or unphosphorylated Spt5 CTD peptides. We found that the pSpt5 peptide, but not the unphosphorylated peptide nor a phosphorylated Rpb1-CTD peptide, effectively competed for binding to ssDNA (Figure 3E, S6D, and S6E). Therefore, pSpt5 and nucleic acid interact with the Plus3 domain on overlapping binding surfaces in a mutually exclusive manner.

To determine the physiological relevance of nucleic acid binding, we conducted *in vivo* examination of R262E and R296E mutants. Both the *prf1-R262E* and *prf1-R296E* mutations significantly decreased recruitment of Prf1 to chromatin in ChIP-qPCR assays; *prf1-R262E* conferred a ~5-fold decrease similar to *prf1-R227A*, whereas *prf1-R296E* conferred a more modest ~2-fold decrease (Figure 4A and S7). Neither of the *prf1-R262E* and *prf1-R296E* mutations significantly affected Prf1 protein or H2Bub1 levels (Figures 4B, 4C, and S3). These data define nucleic acid binding as a biochemical activity distinct from pSpt5 binding that is required for Prf1 association with transcribed genes.

**Figure 4.**
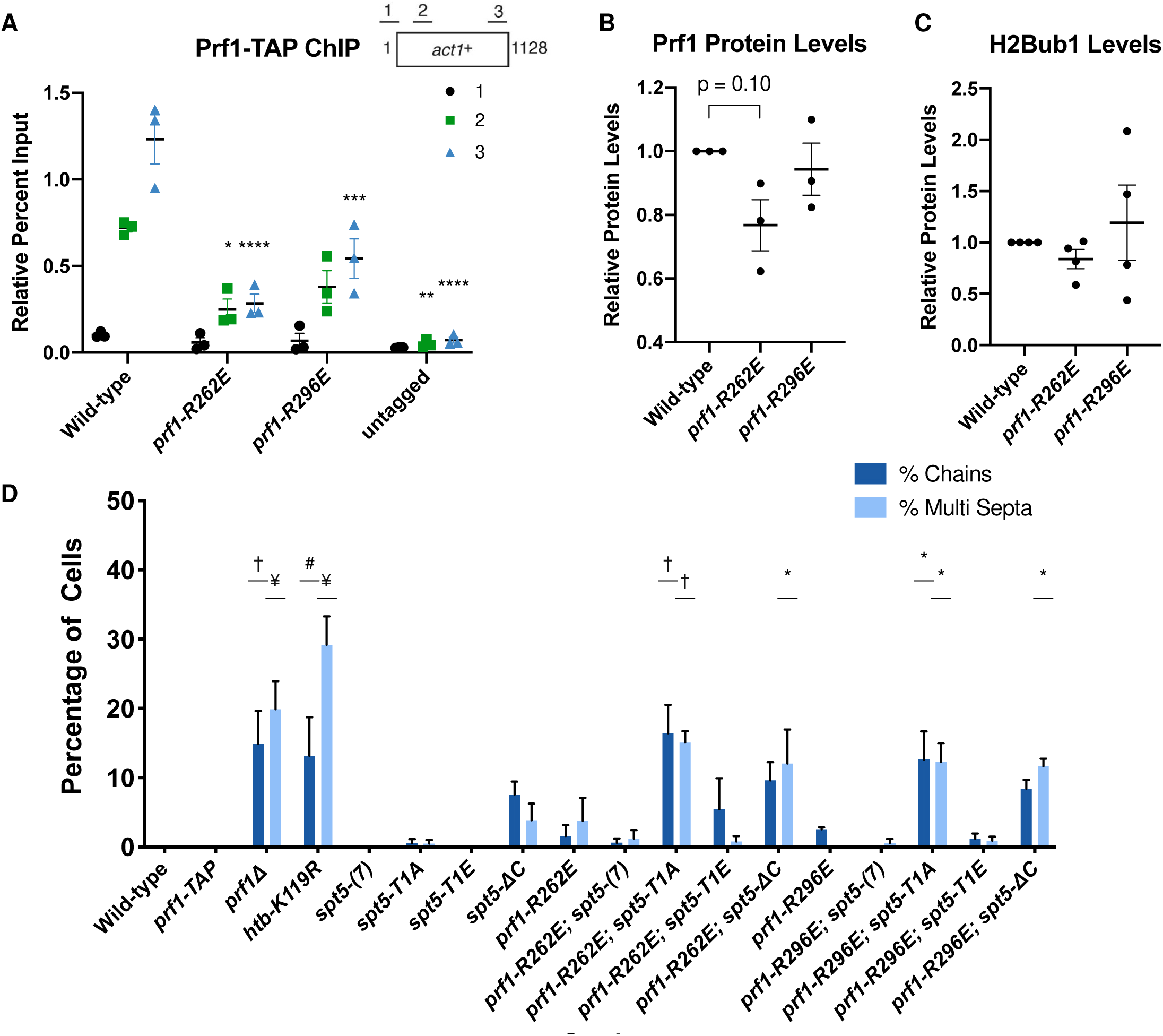
Disruption of Plus3 domain nucleic acid binding and pSpt5 binding have similar phenotypic outcomes. **(A)** TAP-tag ChIP was performed on the indicated strains and quantified with qPCR using the indicated primers in *act1*^*+*^; % IP values were normalized using a primer pair in the *S. cerevisiae PMA1* gene. Length of gene (in base pairs) and position of PCR amplicons shown in diagram at the top. Error bars denote standard error of the mean from 3 independent experiments. A two-way ANOVA was conducted followed by two-sided t-tests with Bonferroni correction between each strain and wild-type within a specific primer pair. * p≤ 0.05, ** p≤ 0.01, *** p≤ 0.001, **** p≤ 0.0001. **(B)** Quantification of immunoblots analyzing Prf1-TAP protein levels normalized to tubulin and then wild-type of the indicated strains. **(C)** Quantifications of H2Bub1 levels normalized to total H3 levels and then wild-type of the indicated strains. For (B) and (C), error bars denote standard error of the mean from 3 independent experiments. A one-sample two-sided t-test was conducted between each strain and its relative normalized wild-type. **(D)** Quantification of septation defects in indicated strains normalized to the number of septated cells counted in each indicated strain. At least 100 cells were counted for each strain per experiment. Error bars denote standard error of the mean from 3 independent experiments. A one-way ANOVA was conducted across all strains followed by two-sided t-tests with Bonferroni correction between each strain and the wild-type *prf1-TAP* strain, for each specific morphology defect. * p≤ 0.05, # p≤ 0.01, † p≤ 0.001, ¥ p≤ 0.0001.

The *prf1-R262E* and *prf1-R296E* mutants displayed phenotypic profiles that were very similar to that of *prf1-R227A*: they did not show any cell division/morphology deficits or drug sensitivity on their own, but showed strong synthetic phenotypes in combination with *spt5-T1A* (Figure 4D and S8). The *prf1-R296E* mutation had milder synthetic effects with *spt5-T1A* on drug sensitivity than either of the other Plus3 domain mutations, although it interacted strongly with *spt5-ΔC* (Figure S8). This may be a reflection of its milder effect on Prf1 function in the ChIP assay (Figure 4A). Prf1 protein levels in the *prf1-R262E spt5* and *prf1-R296E spt5* double mutants were not significantly different in comparison to Prf1 levels from the respective *spt5* single mutant (Figure S9). These results show that the nucleic acid binding surface of the Plus3 domain is important for *in vivo* function of Prf1 independently of the pSpt5 interaction.

### Evidence that the Plus3 domain and the Prf1 C-terminus share a common function

In an effort to characterize other functional regions of *S. pombe* Prf1, we analyzed a series of truncations of the Prf1 C-terminus, a region of the protein previously implicated in binding to the PAF complex (17,24,25). C-terminal truncation mutants terminating at amino acids 345, 458, or 472 (*prf1-Δ345*, *prf1-Δ458*, *prf1-Δ472*) were still recruited to transcribed genes by ChIP-qPCR at levels similar to or even greater than those for wild-type (Figure 5A and S10). The increased ChIP-qPCR signals correlated with increases in Prf1 protein levels by immunoblot (Figures 5B and S3). However, H2Bub1 levels were decreased in all three mutants (Figure 5C). Thus, C-terminally truncated Prf1 proteins are functionally impaired in a manner distinct from Prf1 mutants that disrupt the Plus3 domain.

**Figure 5.**
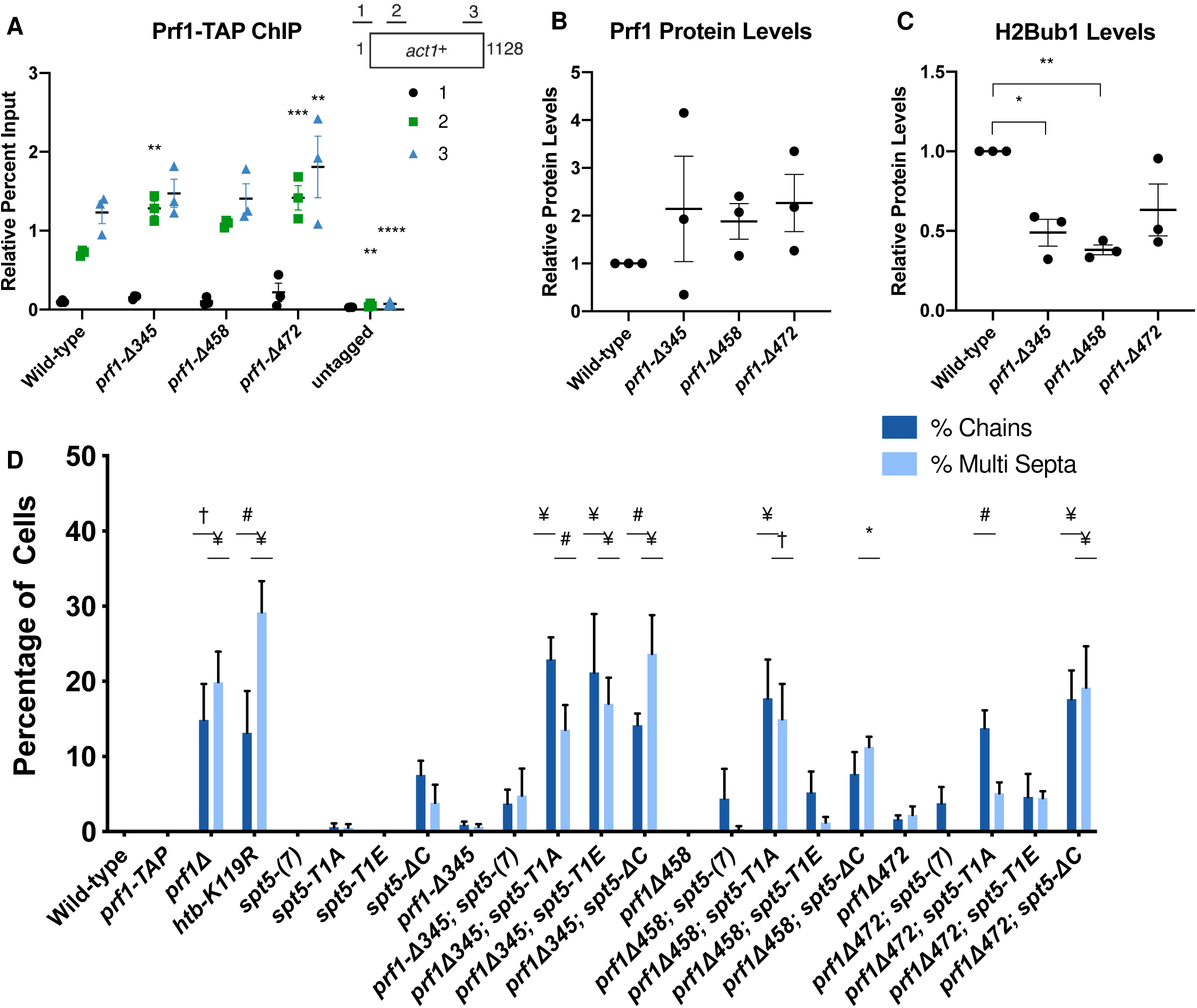
The Prf1 C-terminal region and the Plus3 domain have a shared function. **(A)** TAP-tag ChIP was performed on the indicated strains and quantified with qPCR using the indicated primers in *act1*^*+*^; % IP values were normalized using a primer pair in the *S. cerevisiae PMA1* gene. Length of gene (in base pairs) and position of PCR amplicons shown in diagram at the top. Error bars denote standard error of the mean from 3 independent experiments. A two-way ANOVA was conducted followed by two-sided t-tests with Bonferroni correction between each strain and wild-type within a specific primer pair. ** p≤ 0.01, *** p≤ 0.001, **** p≤ 0.0001. **(B)** Quantification of immunoblots analyzing Prf1-TAP protein levels normalized to tubulin and then wild-type of the indicated strains. **(C)** Quantifications of H2Bub1 levels normalized to total H3 levels and then wild-type of the indicated strains. For (B) and **(C)**, error bars denote standard error of the mean from 3 independent experiments. A one-sample two-sided t-test was conducted between each strain and its relative normalized wild-type. * p≤ 0.05, ** p≤ 0.01. **(D)** Quantification of septation defects in indicated strains normalized to the number of septated cells counted in each strain. At least 100 cells were counted for each strain per experiment. Error bars denote standard error of the mean from 3 independent experiments. A one-way ANOVA was conducted across all strains followed by two-sided t-tests with Bonferroni correction between each strain and the wild-type *prf1-TAP* strain, for each specific morphology defect. * p≤ 0.05, # p≤ 0.01, † p≤ 0.001, ¥ p≤ 0.0001.

The Prf1 C-terminal truncations did not give rise to cell growth and morphology phenotypes on their own (Figure 5D). However, like Plus3 domain mutations, they exhibited synthetic phenotypes in combination with mutations in the Spt5 CTD. For the *prf1-Δ472* and *prf1-Δ458* double mutant strains, the synthetic phenotypes were observed for all assays tested in combination with *spt5-ΔC.* Double mutants with *spt5-T1A* exhibited assay-dependent effects: *prf1-Δ472* caused septation defects, and MMS sensitivity, whereas *prf1-Δ458* caused septation defects, TBZ sensitivity, and MMS sensitivity (Figure 5D and S11). Modest septation phenotypes were observed in double mutants with *spt5-T1E* but these were not statistically significant, suggesting that the function of this C-terminal portion of Prf1 is related to loss of Spt5 CTD phosphorylation (Figure 5D and S11). The largest *prf1* truncation mutation, *prf1-Δ345*, displayed cell division/ morphology deficits and drug sensitivity with the *spt5-T1A*, *T1E* and Δ*C* mutants (Figure 5D and S11). The fact that alanine and glutamate substitutions at the T1 position were similarly deleterious in this background suggests that the larger truncation impinges on a function that is either stringently dependent on T1, or dependent on phosphorylated T1 in a way that is not compensated by the negatively charged side-chain. Levels of the C-terminally truncated Prf1 proteins were unchanged or increased compared to those of the wild-type Prf1 in the respective *spt5* mutant backgrounds (Figure S12). Taken together, these results suggest that amino acids 459-562 of Prf1 participate in an interaction that functions in parallel with Spt5-T1 phosphorylation, similar to the Plus3 domain, and that amino acids 345-458 of Prf1 participate in an additional function that is more generally sensitive to Spt5 CTD structure.

### Prf1 Plus3 domain and C-terminal region both interact with the PAF complex

We hypothesized that the Prf1 Plus3 domain and C-terminal region may share a common physical interactor that accounts for their shared function. This is unlikely to be nucleic acid, as we have not detected any nucleic acid binding by the Prf1 C-terminal region (data not shown). The C-terminal region also has no affinity for the Spt5 CTD (Figure S1). Given that the PAF complex has previously been shown to interact with the C-terminal regions of human and *S. cerevisiae* Rtf1, we investigated interaction between Prf1 and PAF complex using purified proteins (17,24,25). We observed that full-length Prf1, the Plus3 domain, and the C-terminal region (amino acids 345-562), produced as recombinant GST fusion proteins (Figure S1A), associated with native *S. pombe* PAF complex (purified via the TAP method; Figure 6A) in GST-pulldown experiments (Figure 6B). The N-terminal HMD domain did not pull down PAF, indicating that PAF interacts specifically with the Plus3 domain and C-terminal region *in vitro* (Figure 6B). As interaction between the Plus3 domain and PAF has not previously reported, we used surface plasmon resonance (SPR) to confirm it. Specific dose-dependent binding between the PAF complex and the Plus3 domain was also apparent in SPR experiments (Figure S13A). Importantly, the R227A, R262E, and R296E mutations in the Plus3 domain all reduced this interaction (Figure 6C, Figure S13B). As these mutations all affect nucleic acid binding with the Plus3 domain, we tested whether the interaction between the Plus3 domain and PAF is nucleic acid-dependent. We observed similar interaction between Plus3 domain and PAF in the presence of ethidium bromide, suggesting that it reflects a direct protein-protein interaction (Figure S13C). Indeed, addition of exogenous ssDNA reduced the efficiency of the Plus3 domain-PAF interaction in GST pulldowns (data not shown). Thus, interaction with the PAF complex is an additional Plus3 domain function that may operate in parallel with pSpt5. We have not yet identified genetic interactions between PAF complex mutations and *spt5* CTD mutations that support this (data not shown). This is likely due to the fact that interaction of the Plus3 domain with the PAF complex seems to involve multiple individual PAF complex subunits based on our *in vitro* characterization, including Leo1, the N-terminal half of Tpr1 (Tpr1N), and the C-terminal half of Paf1 (Paf1C) (Figure S13D-H). However, ChIP-qPCR assays showed a significant decrease in *prf1-TAP* chromatin occupancy at the *act1*^*+*^, *spbc354.10*^*+*^ and *nup189*^*+*^ genes in a *tpr1Δ* strain compared to wild-type (Figure 6D-F). As *tpr1Δ* is predicted to eliminate the PAF complex, this indicated that PAF, like the Plus3 domain and pSpt5, is necessary for Prf1 chromatin association. PAF chromatin occupancy showed a locus-specific dependence on Prf1, as *paf1-TAP* recruitment was affected by *prf1Δ* at *act1*^*+*^ but not at the other two loci; this may reflect locus-specific functions for the Prf1-PAF interaction (Figure 6F). These data suggest that a direct Prf1-PAF interaction, mediated in part by the Plus3 domain, promotes Prf1 function in conjunction with pSpt5.

**Figure 6.**
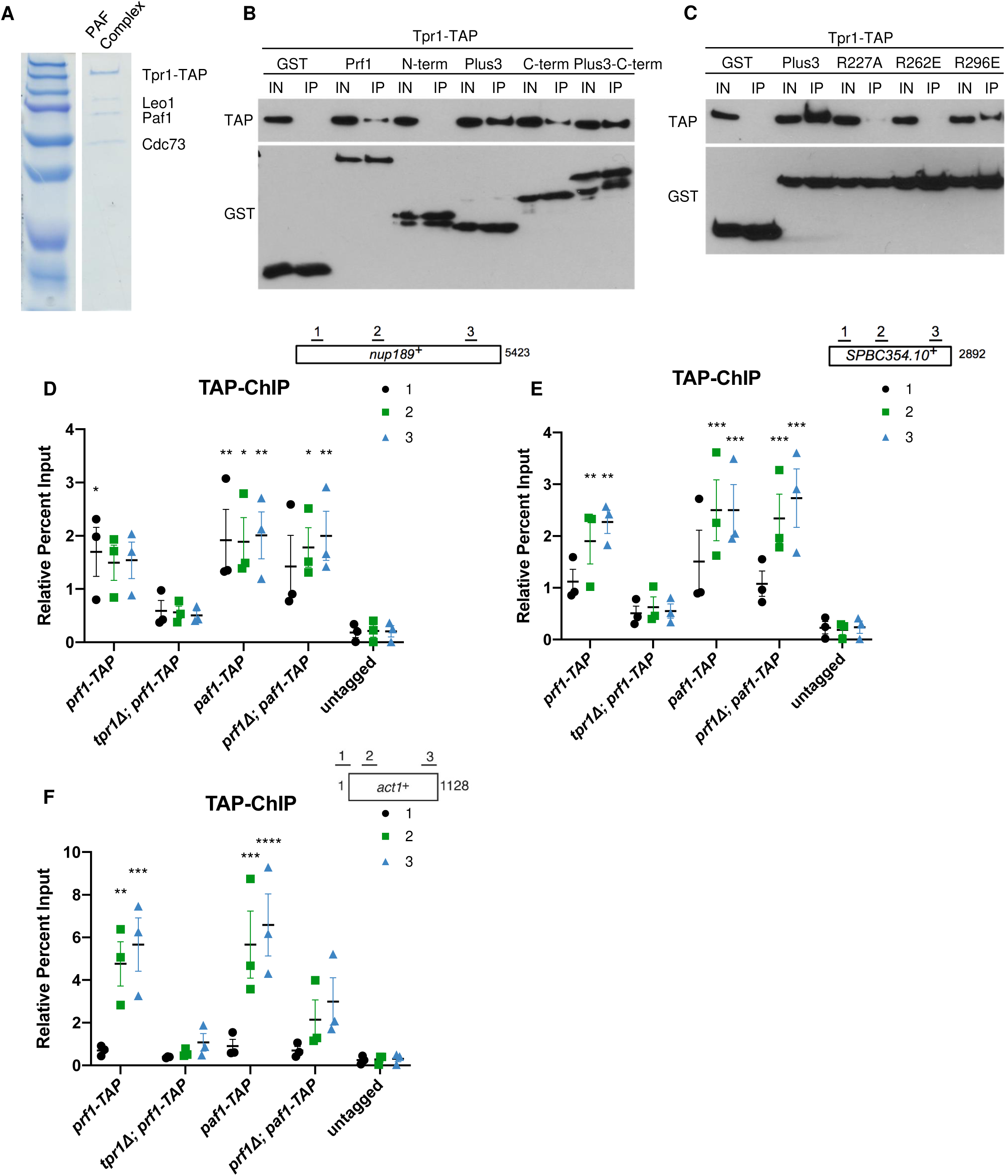
Prf1 interacts with the PAF Complex through its Plus3 domain and C-terminal region. **(A)** Native PAF complex purified from a *tpr1-TAP* strain was analyzed by SDS-PAGE and Coomassie staining. Subunits of the complex are labeled on the right; size markers are indicated on the left. “Tpr1-CBP” refers to Tpr1 fused to calmodulin-binding peptide that is present after TAP purification. **(B)** GST pulldowns of the native PAF Complex (purified via Tpr1-TAP) with full-length, recombinant GST-Prf1 or the indicated domains tagged with GST were analyzed by SDS-PAGE and immunoblotting with the indicated antibodies (left). “Plus3-C-term” denotes a fragment of Prf1 consisting of both the Plus3 domain and C-terminus. “I” denotes input (5%); “IP” denotes bound fraction (50%). All experiments were repeated at least 3 independent times and representative blots are shown. **(C)** As in (B) with the indicated GST-Plus3 domain fusions. **(D-F)** TAP-tag ChIP was performed on the indicated strains and quantified with qPCR using the indicated primers in *nup189*^*+*^, *spb354*^*+*^, and *act1*^*+*^; % IP values were normalized to the input of each corresponding strain and primer pair. Length of gene (in base pairs) and position of PCR amplicons shown in diagram at the top. Error bars denote standard error of the mean from 3 independent experiments. A two-way ANOVA was conducted followed by two-sided t-tests with Bonferroni correction between each strain and untagged within a specific primer pair. * p≤ 0.05, ** p≤ 0.01, *** p≤ 0.001, **** p≤ 0.0001.

## Discussion

This study provides novel insights into the function of the Rtf1 Plus3 domain and its relationship to the Spt5 CTD. Previous studies have centered on the direct interaction between the Plus3 domain and Spt5 CTD repeats phosphorylated at the conserved T1 position and have emphasized its importance for recruitment of Rtf1 and the PAF complex to transcribed genes (14,15). Our genetic and biochemical analyses strongly argue that 1) the Plus3 domain engages in an additional interaction, exclusive of that with pSpt5, that is critical for Prf1/Rtf1 function and recruitment *in vivo*; and 2) pSpt5 promotes Prf1/Rtf1 function in parallel through another factor.

We observed broad phenotypic overlap between *prf1-R227A*, which abolishes pSpt5 recognition, and both *prf1-R262E* and *prf1-R296E*, which retain pSpt5 binding. The phenotypic effects of these mutations were strongest in *spt5-T1A* and *spt5-ΔC* genetic backgrounds, and absent or weak in combination with *spt5-T1E*, indicating that introduction of a negative charge at the T1 position is important for Prf1 function when the Plus3 domain is compromised. The fact that phenotypic enhancement was observed with *spt5-ΔC* (albeit to varying extents) negates the possibility that another CTD phosphorylation site is bound by the mutant Plus3 domains, or that Plus3 domain binding to the unmodified CTD drives the phenotypic effects. We cannot exclude the possibility that a physical interaction occurs between Prf1 and Spt5 that is independent of the Spt5 CTD altogether but that requires the Plus3 domain. This would be an entirely different interaction than that suggested by previous work in budding yeast (14).

All of these mutations reduce Prf1 occupancy on gene coding regions by ChIP. This effect is particularly pronounced for *prf1-R227A* and *prf1-R262E*, both of which exhibit occupancy levels close to background. Thus, the pSpt5-independent interaction of the Plus3 domain is important for Prf1 recruitment to chromatin, consistent with the role of the Plus3 domain previously defined in *S. cerevisiae* (14,17). The *spt5-T1A* mutation reduces Prf1 chromatin occupancy as well as Prf1 protein levels, although we argue that these effects are not solely attributable to interaction with Prf1 (16). Thus, Prf1 chromatin occupancy requires both Plus3 domain function and pSpt5, but Prf1 function can be maintained in the absence of either one. These findings suggest that Prf1 function does not require its stable association with chromatin and is compatible with more dynamic associations that are not captured by ChIP (42).

Recombinant Plus3 domain binds to purified, native PAF complex in a manner that is also sensitive to *prf1-R227A, prf1-R262E*, and *prf1-R296E* mutations. Prf1 interaction with the PAF complex also involves its C-terminal region, truncation of which leads to synthetic phenotypes in combination with *spt5-T1A*. Direct interaction between Prf1 and PAF was previously demonstrated using purified components and was primarily attributed to the C-terminal region of Prf1 (25,43). Our finding that PAF can also directly interact with the Plus3 domain is also consistent with crosslinking mass spectrometry analysis of the *S. cerevisiae* Rtf1/PAF complex, in which both Plus3 and C-terminal regions could be crosslinked to PAF (43). These results support a model in which the Plus3 domain and the C-terminal region both interact with the PAF complex, thereby promoting Prf1 function in parallel to pSpt5. This idea is further supported by the fact that the PAF complex is necessary for Prf1 chromatin occupancy. However, given that Prf1 function is maintained in cases where its chromatin occupancy is greatly reduced, how interaction with PAF contributes to Prf1 function remains unclear. Greater insight into the significance of this interaction will require identification of additional pSpt5-binding factors, as the genetics argues that pSpt5 contributes in parallel to PAF’s function in this context. We detect interaction between multiple PAF subunits and the Plus3 domain *in vitro*, but interactions between Prf1 and the PAF complex are weak or undetectable in extracts (as is the case in metazoans), further complicating efforts to dissect the function of the interaction *in vivo* (16,44). A more detailed picture of the molecular basis for the cell division and morphology phenotypes of *prf1* will also be critical to understanding the significance of Prf1 interactions.

The *prf1-R227A, prf1-R262E*, and *prf1-R296E* mutations all impair a nucleic acid binding activity of the Plus3 domain. This activity prefers ssDNA over dsDNA, as has been demonstrated previously for the human Plus3 domain (18). We also show that the affinity of Prf1 Plus3 domain for RNA is similar to that for ssDNA. This is consistent with studies showing interaction of *S. cerevisiae* Rtf1 with RNA *in vitro* and *in vivo* (45,46). Whether or not nucleic acid is a physiologically relevant binding partner for the Plus3 domain *in vivo* remains to be determined. It is clear, however, that the binding of the Plus3 domain to pSpt5 and nucleic acid are mutually exclusive, because 1) *prf1-R227A* abrogates both, and 2) pSpt5 competes with nucleic acid for Plus3 domain binding. Nucleic acid also competes with PAF complex for Plus3 binding (data not shown). The differential effects of R262E and R296E on nucleic acid binding (and PAF complex binding) versus pSpt5 binding suggest that the interaction interface for the former may be larger. Nonetheless, results of the competition experiments suggest that the Plus3 domain can interact with multiple partners through a common interface (or distinct but overlapping interfaces). We suggest that multiple Plus3 domain interactions could occur in the context of an extended Spt5 CTD with multiple phosphorylated repeats. Whereas Prf1 may directly bind to pSpt5 at some repeats, alternate modes of association may predominate at others. Our data also suggest that all modes of interaction are needed to observe stable association of Prf1 with chromatin, but that this apparent plasticity could explain how function is maintained when either pSpt5 or the Plus3 domain is compromised. Determining whether this plasticity might be regulated, and what the functional consequences might be for transcription, are important avenues for future study.

Our results point to additional Spt5 CTD interactors that are regulated by CTD phosphorylation and that promote function of Prf1/Rtf1. Few direct interactions with the Spt5 CTD have been described previously, and the Plus3/CTD interaction is the only one known to be phospho-specific. Factors involved in 5’ and 3’ mRNA processing also interact with the Spt5 CTD (47–49). Phospho-specificity of cleavage and polyadenylation factor interaction with the Spt5 CTD has not been determined, whereas capping enzyme interaction is blocked by T1 phosphorylation (47). Interestingly, we observed that T1 phosphorylation also blocks interaction of the PAF complex with the Spt5 CTD, although the physiological relevance of this interaction is not known (16). Further investigation of the functional relationship between the Spt5 CTD and Prf1/Rtf1 may uncover novel mechanisms linking Spt5 CTD phosphorylation to RNAPII elongation control.

## Acknowledgements

We thank K. Gull for kindly providing TAT1 monoclonal antibody against alpha-tubulin; R. Fisher, B. Schwer and S. Shuman for *S. pombe* strains; F. Robert for *S. cerevisiae* expressing *RPB1-TAP*; and members of the Tanny lab for helpful discussions. This work was supported by the Canadian Institutes for Health Research (MOP-130362 to J.C.T., MOP-142184 to D.C.), and Natural Sciences and Engineering Research Council of Canada (RGPIN 03661-15 to J.C.T., RGPIN-2015-04848 to D.C.). J.C.T. acknowledges support from the Fonds de recherche santé Quebec (chercheur boursier 33115). The McGill SPR-MS Facility is supported by the Canada Foundation for Innovation.

